# Dual RNAseq shows the human mucosal immunity protein, MUC13, is a hallmark of *Plasmodium* exoerythrocytic infection

**DOI:** 10.1101/183764

**Authors:** Gregory M. LaMonte, Pamela Orjuela-Sanchez, Lawrence Wang, Shangzhong Li, Justine Swann, Annie N. Cowell, Bing Yu Zou, Alyaa M. Abdel-Haleem Mohamed, Zaira Villa-Galarce, Marta Moreno, Carlos Tong-Rios, Joseph M. Vinetz, Nathan Lewis, Elizabeth A. Winzeler

## Abstract

The exoerythrocytic stage of *Plasmodium* malaria infection is a critical window for prophylactic intervention. Using a genome-wide dual RNA sequencing of flow-sorted infected and uninfected hepatoma cells we identify the human mucosal immunity gene, *Mucin13 (MUC13)*, as strongly upregulated during *Plasmodium* exoerythrocytic hepatic-stage infection. We confirm that *MUC13* expression is upregulated in hepatoma cell lines and primary hepatocytes. In immunofluorescence assays, host MUC13 protein expression distinguishes infected cells from adjacent uninfected cells and shows similar colocalization with parasite biomarkers such as UIS4 and HSP70. We further show that localization patterns are species independent, distinguishing both *P. berghei* and *P. vivax* infected cells, and that MUC13 can be used to identify compounds that inhibit parasite replication in hepatocytes across all Human-infecting *Plasmodium* species. This data presents a novel interface of host-parasite interactions in *Plasmodium*, in that a component of host mucosal immunity is reprogrammed to assist the progression of infection.

## Introduction

Malaria remains a significant global health problem with 214 million annual cases and up to a half million deaths in 2015 (W.H.O., 2017). The disease, caused by protozoan parasites of the genus *Plasmodium*, is transmitted when a female *Anopheline* mosquito takes a blood meal and injects infectious *Plasmodium* sporozoites. These sporozoites (typically less than 100) migrate to the liver where they invade hepatocytes. This exoerythrocytic infection develops asymptomatically in the infected hepatocytes over a period of two to ten days, depending on the species of malaria parasite. The infected hepatocyte eventually bursts, releasing tens of thousands of merozoites that are programmed to infect erythrocytes. The repeated infection and lysis of erythrocytes results in symptomatic disease, and for this reason, the erythrocytic stage has been the historical focus of drug discovery. On the other hand, the exoerythrocytic stage attracts attention due to the substantially reduced parasite burden. Unsurprisingly then, most malaria vaccine candidates (such as RTS,S (Olotu et al., 2016), also known as Mosquirix) target the exoerythrocytic stage for this reason. In addition, while malaria is typically prevented through the use of insecticide-treated bed nets and treated with chemotherapy such as artemisinin combination therapies, there is a recognized need for new molecules that may protect against malaria and which might be formulated as a component in a SERCaP (Single Exposure, Radical Cure, and Prophylaxis) medicine that could be used in a malaria-elimination campaign (Burrows et al., 2013).

From the perspective of host-parasite interactions, there are likely numerous possible host targets for therapeutic intervention. During the initial stage, the infected hepatocyte can grow to many times its initial size and yet does not undergo apoptosis. The parasite’s metabolic needs are also likely to be considerable given that one sporozoite can yield over 30,000 merozoites within a single infected host cell. It thus seems very likely that the parasite is releasing effectors into the host cell to control host cell behavior. This notion that the malaria parasite is modifying host-gene expression is heavily supported by studies in the related apicomplexan parasite, *Toxoplasma gondii*. Gene expression studies in *T. gondii* have been used effectively to characterize the host response to infection, due to its high multiplicity of infection (Melo et al., 2013; Saeij et al., 2007). As observed in these studies, the parasite must carefully regulate immune activation and host-cell effector mechanisms (reviewed in (Melo et al., 2011)) to establish infection. It is now known that multiple proteins, including ROP18 kinase (Saeij et al., 2006; Taylor et al., 2006) and GRA15 (Rosowski et al., 2011), are secreted into the host cell, altering host cell signal transduction and inflammation (Jensen et al., 2013).

In contrast to *Toxoplasma*, relatively little is known about molecules produced by hepatocytes in response to *Plasmodium* sporozoite infection, in part because of the difficulty associated with studying the exoerythrocytic stage (reviewed in (Ingmundson et al., 2014)). *Plasmodium* sporozoites form a parasitophorous vacuole within infected hepatocytes. Parasite-induced host molecules are known to include EphA2 and CD81, which have been shown to be essential for hepatocyte invasion (Kaushansky et al., 2015; Yalaoui et al., 2008). Parasite-secreted molecules include LISP and IBIS1, which are secreted into hepatocytes in the *P. berghei* model (Ingmundson et al., 2012; Orito et al., 2013). Another candidate effector molecule is the circumsporozoite protein (CSP), an 3 abundant protein that is shed from the parasite sporozoite surface. It was also shown that expression of recombinant CSP in HeLa cells regulates TNF-alpha dependent host-immune signaling and NF-?B translocation to the nucleus, for example (Singh et al., 2007).

As with *Toxoplasma*, global gene expression analysis of infected cells should be an effective way to identify host genes that play a role in *Plasmodium* exoerythrocytic infection. However, the low parasite to hepatocyte ratio also creates a low signal to noise ratio. This problem can be overcome using dual-RNA sequencing of flow-sorted infected host cells (Cloney, 2016), which analyzes host and pathogen transcriptomes simultaneously. In addition, the tremendous depth of coverage offered by current transcriptomic sequencing approaches allows for a deep examination of the data. Here, we have taken advantage of this dual-RNA sequencing approach in order to gain insights into *Plasmodium* liver-stage development. We have identified and partially characterized one host-factor, *Mucin13*, which is highly upregulated during host-cell infection by both *P. berghei* and *P. vivax*. In contrast to previously identified parasite markers of liver-stage infection, such as HSP70 or UIS4 (Matuschewski et al., 2002). Mucins, as a component of the mucosal immune system, serve a critical role in host innate immune defenses as an initial barrier to infection. MUC13, as one of the transmembrane mucins, has been shown to play important roles in defense against several forms of cancer (Sheng et al., 2017) and bacterial infection, such as by *Helicobacter pylori* (Liu et al., 2014). Mucosal immunity has never been shown to have a significant role in malaria (reviewed in (Gazzinelli et al., 2014)), in part due to the lifecycle of the parasite itself. However, in this study, we display a novel interaction of innate immunity against parasite infection, in which activation of MUC13 aids parasite development. Therefore, MUC13 may serve not only as a universal biomarker for *Plasmodium* infection, upregulated regardless of the species of infecting malaria parasite, but also plays a role in parasite development as well.

## Results

To investigate host-pathogen interactions in *Plasmodium* exoerythrocytic stages, we conducted a dual RNA sequencing study, a strategy that has proven useful in identifying novel interactions in other intracellular parasitic organisms, including *T. gondii* and *Leishmania major* (Dillon et al., 2015; Pittman et al., 2014; Westermann et al., 2017). We initially used *P. berghei*, a cause of murine malaria to infect the hepatoma cell line Huh7.5.1, which had proven to be readily sortable using flow cytometry (Swann et al., 2016). The Huh7.5.1 hepatocytes were first infected *in vitro* with *P. berghei* sporozoites expressing GFP regulated by the elongation factor-1α promoter (Franke-Fayard et al., 2004) that is active throughout the 48-hour liver stage infection (Swann et al., 2016). Samples were collected at time 0 (uninfected hepatocytes and sporozoites before infection), 24 hours, and 48 hours post-infection (hpi), then infected and uninfected cells were separated using fluorescence-activated cell sorting. Altogether, approximately 10,000 cells of each type were collected, and based upon a pilot study we determined that obtaining 20 million reads at 24 hours post-infection, or 10 million reads at 48 hours post-infection, would be sufficient to obtain an average 50X coding genome coverage in both host and *P. berghei* (Figure S1). Huh7.5.1 experiments were conducted in triplicate on three different days. To ensure that our expression patterns were not dependent on the human cell line used, we collected additional experiments with HepG2-CD81 cells (48 hours infected and uninfected) and twice with HC04 cells (48 hours infected and uninfected), both additional human hepatocyte cell lines able to support *P. berghei* infection (Sinnis et al., 2013). Total RNA was isolated from the different populations and dual RNA-sequencing performed for both *P. berghei* and the host human hepatocyte cell lines at each time point. Over 318 million total reads were obtained and aligned to either the *P. berghei* reference genome or the human reference genome, 82.1% of which map to either the human of *P. berghei* genomes.

Altogether we detected transcripts with at least one sample > 10 reads for 21,941 of the 58,051 human genes and 4770 of the 5245 parasite genes (Table S1, S2). Expression patterns were generally consistent across experimental replicates (R^2^= 0.86 to 0.89). In the parasite, PBANKA_050120 (*UIS4*), and PBANKA_040320 (*CSP*), were the most highly expressed at time zero. PBANKA_071190 (*HSP70*), as well as PBANKA_100300 (*LISP2*), were the most highly expressed at 48 hours (Table S2). All are well validated hepatic stage markers (Orito et al., 2013; Vaughan et al., 2012).

We next identified genes that were differentially expressed during infection. While our ultimate goal was to identify host biomarkers of infection, we first analyzed the gene expression patterns for *P. berghei* as a control to ensure that our time point accurately reflected the expected parasite biology. As anticipated, parasite gene expression changes followed known patterns, including downregulation of genes with known sporozoite function, such as *CSP* (log_2_FC of −9.6, p-value 3.61 × 10^−109^), *CELTOS* (log_2_FC of −12.3, p-value 3.70 × 10^−78^), and *UIS4* (log_2_FC of −9.3, p-value 9.32 × 10^−219^), combined with upregulation of merozoite genes such as *MSP1* (log^2^FC of 3.8, p-value 2.78 × 10^−25^), *MSRP2* (log_2_FC of 6.6, p-value 8.22 × 10^−23^) and *SERA1* (log_2_FC of 5.0, p-value 2.84 × 10^−46^) (Table S2) (Tarun et al., 2008). We also observed excellent concordance between our *P. berghei* expression data and reported *P. falciparum* quantitative microarray gene expression studies, with the *P. falciparum* homolog of 5 of the 10 most highly expressed *P. berghei* sporozoite transcripts (by read count at 0 hpi) also being among the 10 most highly expressed *P. falciparum* transcripts (*CSP, ETRAMP10.3/UIS4, TRAP, CELTOS and HSP70* - hypergeometric mean probability of association by chance < 1.51 × 10^−12^) (Le Roch et al., 2003). In addition, 13 of the 20 most highly expressed *P. falciparum* sporozoite transcripts (Le Roch et al., 2003) possessed homologs that were also highly expressed (within the top 100 genes) in our *P. berghei* dataset (p< 2.82 × 10^−18^), demonstrating that the sporozoite transcriptional profile is largely shared between rodent and human *Plasmodium* species regardless of the experimental platform.

After having demonstrated the validity of our experimental design, we turned our focus to our primary goal, the identification of host-factors involved in parasite development. In order to broadly examine host-cell gene expression patterns, we first identified differentially expressed genes using pairwise comparisons between infected and uninfected cells focusing primarily on the 48-hour time point (which was sampled with multiple hepatocyte lines). Genes were next subjected to hierarchical clustering (Figure 1A, Matrix used to create tree - Table S3). The data showed, not unexpectedly, that there were major differences between the different cell lines. However, subclusters could be identified that revealed genes that were consistently upregulated across cell lines with the most dramatic changes occurring 48 hpi.

**Figure 1:**
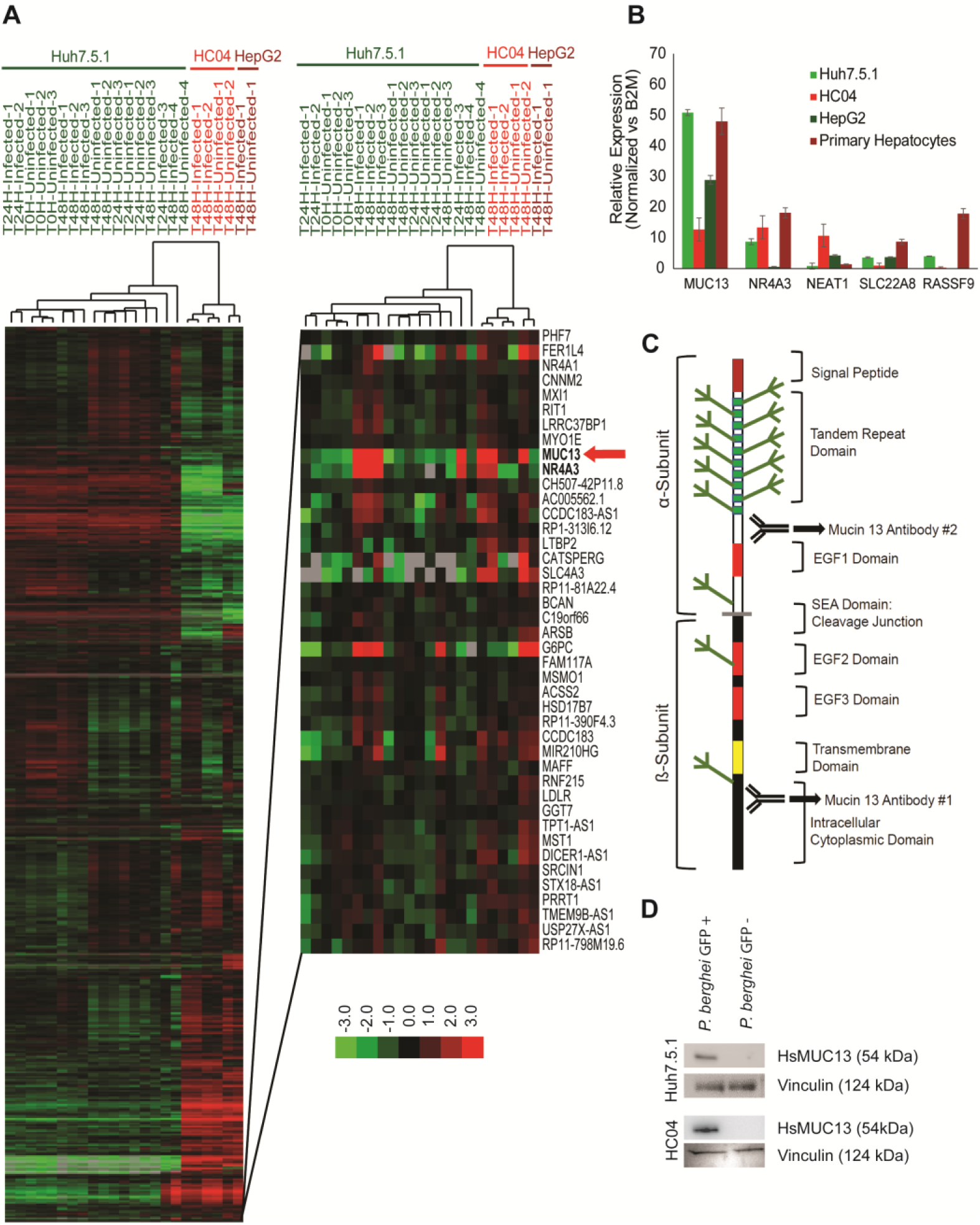
Identification of differentially expressed host and parasite transcripts. (A) Hierarchical Clustering of RNAseq samples based on gene expression patterns in the indicated host hepatocytes and specified times post-infection. (B) Confirmation of host factor upregulation via qPCR in the indicated hepatocyte cell types. (C) Schematic of MUC13 protein structure and approximate antibody location. (D) Confirmation of *MUC13* upregulation during infection via western blot.

In order to classify the gene expression changes, we determined whether specific functional classes were overrepresented in up and down regulated genes. Analysis of the 21,941 detected genes at the 48-hour time point showed that 840 human genes were upregulated (average >2X change between pairs, p <0.01) and 618 were downregulated using similar criteria. The sets of upregulated genes were compared to compiled lists of genes with known functions, including gene ontology groups using Metascape (Tripathi et al., 2015). We found that genes involved in host ribosome function (p = 3.23 × 10^−14^), and DNA replication (p = 3.46 × 10^−10^) were downregulated (Table S4), indicating that the host cell has decreased the production of ribosomes and DNA replication machinery as it is presumably no longer dividing. The most strongly downregulated gene, *ENHO*, plays a role in the positive regulation of Notch signaling (Energy Homeostasis related, 4.34X average downregulation, p = 1.11 × 10^−08^). The cheomokine ligand, CXCL10, which is a hallmark protein in the JAK-STAT, and alpha and gamma interferon pathways, was also highly downregulated (2.88X downregulation p = 4.68 × 10^−05^), as was thymidine kinase, a gene involved in DNA synthesis. Further analysis of dysregulated transcriptional pathways using Ingenuity pathway analysis (Kramer et al., 2014) also indicated downregulation of the eIF2 signaling (Figure S2).

The 840 upregulated genes were more difficult to classify. Transcript increases were observed for genes that play a role in general transcriptional pathways (Generic Transcription Pathway - p = 2.0 × 10^−6^), as well as genes with a role in the regulation of energy homeostasis (p = 1.32 × 10^−4^) with enrichment in adipocytokine signaling pathway and gluconeogenesis, among others (Table S5). These changes are presumably a reflection of the metabolic changes the host cell needs to make to support rapid parasite growth and replication. We also observe strong upregulation of mTOR signaling, whose upregulation has been shown to be necessary for hepatic development (Hanson et al., 2013), using Ingenuity pathway analysis. The most highly upregulated gene, however, was the human mucosal innate immunity gene, *Mucin*13 (*MUC13,* average 16.51 upregulation, p = 7.84 × 10^−51^), a gene which is also upregulated at the mRNA level in response to *H. pylori* infection or IL-1β stimulation of MKN7 cells (Cheng et al., 2016) or treatment with the colitis-inducing agent, dextran sodium sulfate (Sheng et al., 2011).

Because *Mucin13* (*MUC13*) transcripts are found in some cancers including ovarian and colorectal cancers (Gupta et al., 2014) we next sought to confirm that this was not an artifact of using immortal hepatoma cells. To further ensure that the expression changes were not limited to hepatoma cell lines, we validated upregulation in *P. berghei* infected primary human hepatocytes, as well as in *P. berghei* infected Huh7.5.1, HepG2 and HC04 cells using qRT-PCR. We chose the two clearest examples of infection status marker genes (Figure 1B): *MUC13* and *NR4A3.* In addition, we chose to validate several additional genes (*SLC22A8, RASSF9* and *NEAT1*) which were highly upregulated in Huh7.5.1 cells, yet not within this infection responsive cluster, as controls. We found that in these four cell lines, we observed that only *MUC13 mRNA* was consistently upregulated (∼16X) (Figure 1B).

*MUC13* encodes a membrane bound mucin with abundant O- and N-glycosylation on its extracellular domain (Schematic: Figure 1C) as well as EGF domains and has a molecular weight of 54 kDa. To test whether we also observe upregulation of the MUC13 protein we performed western blot analysis using a commercially available polyclonal antibody that recognizes the C-terminal domain (Figure 1D). Interestingly, MUC13 has been reported both as a 54 kDa transmembrane protein and a smaller 33 kDa (putative) soluble form once the transmembrane domain is cleaved (Maher et al., 2011). We clearly observed the 54 kDa form (Figure 1D), although we did not observe the lower molecular weight product (Figure S3), indicating that the N-terminal extracellular domain may remain attached. In addition to recognizing the correct band, the MUC13 antibody showed protein upregulation only in infected Huh7.5.1 and HC04 cells (Figure 1D), compared to uninfected co-sorted controls that were exposed to identical culture conditions.

To assess whether MUC13 colocalizes with parasite in infected hepatocytes, we used the same antibody described above and contained infected cells using two independent parasite antibodies that react with Heat-shock protein 70 kDa (HSP70) and Upregulated-in-Sporozoites 4 (UIS4) (Figure 2A, B). These two antibodies stain the cytoplasm of developing exoerythrocytic form schizonts (Renia et al., 1990) and the parasite membrane (Mueller et al., 2005) respectively. Our immunofluorescence (IFA) analysis showed that our MUC13 antibody recognized only parasite infected cells. Interestingly, MUC13 is typically expressed on the outer cell membrane of host mucosal epithelial cells (Williams et al., 2001) where it is found on apical membranes and forms part of the glycocalyx that presumably protects gut cells from pathogens. However, in parasite-infected cells the staining (Figure 2A,B) largely colocalized with cytoplasmic parasite HSP70, indicating that this host protein may be actively transported into the parasite cytoplasm. Although the cytoplasmic localization is surprising, it reflects the documented behavior of this protein in cancerous cells. It has been shown that *MUC13* is cytoplasmic and overexpressed in metastatic colon cancer cells but apically located in normal adjacent control cells (Gupta et al., 2012). In addition, MUC13 is translocated from the apical membrane to the cytoplasm in colonic epithelium cells after treatment with the colitis-inducing agent, dextran sodium sulfate (Sheng et al., 2011). While this could be due to acquired cross-reactivity in stimulated cells, MUC13 antibodies reveal no MUC13 staining in knockout animals (Sheng et al., 2011). Although the apparent localization within the parasite cytoplasm could indicate cross-reactivity with parasite proteins, we found that MUC13 IFA levels closely correlate with transcript levels. We observed almost no *MUC13* transcripts in our RNAseq at time zero and an IFA time course also showed no MUC13 staining two hours after infection, during which time parasites were present and could be readily detected by HSP70 antibody staining (Figure 3 and data not shown). The intensity of MUC13 staining was also directly proportional to *MUC13* RNA levels, with a very modest induction at 24 hpi, and increasing to strong induction at 48 hpi (Figure 3). This induction of MUC13 also seems to be specific to exoerythrocytic infection, as IFA analysis for MUC13 in asexual blood stage parasites (with MUC13 Antibody #1 – LifeSpan Biosciences #LS- C345092) shows no colocalization with parasites, only a faint MUC13 signal along the RBC membrane (Figure S4). In addition, staining with an independently derived, commercial rabbit polyclonal antibody to the MUC13 extracellular domain (MUC13 Antibody #2 - LifeSpan BioSciences #LS-A8191) showed nearly identical co-staining patterns (Figure S5).

**Figure 2:**
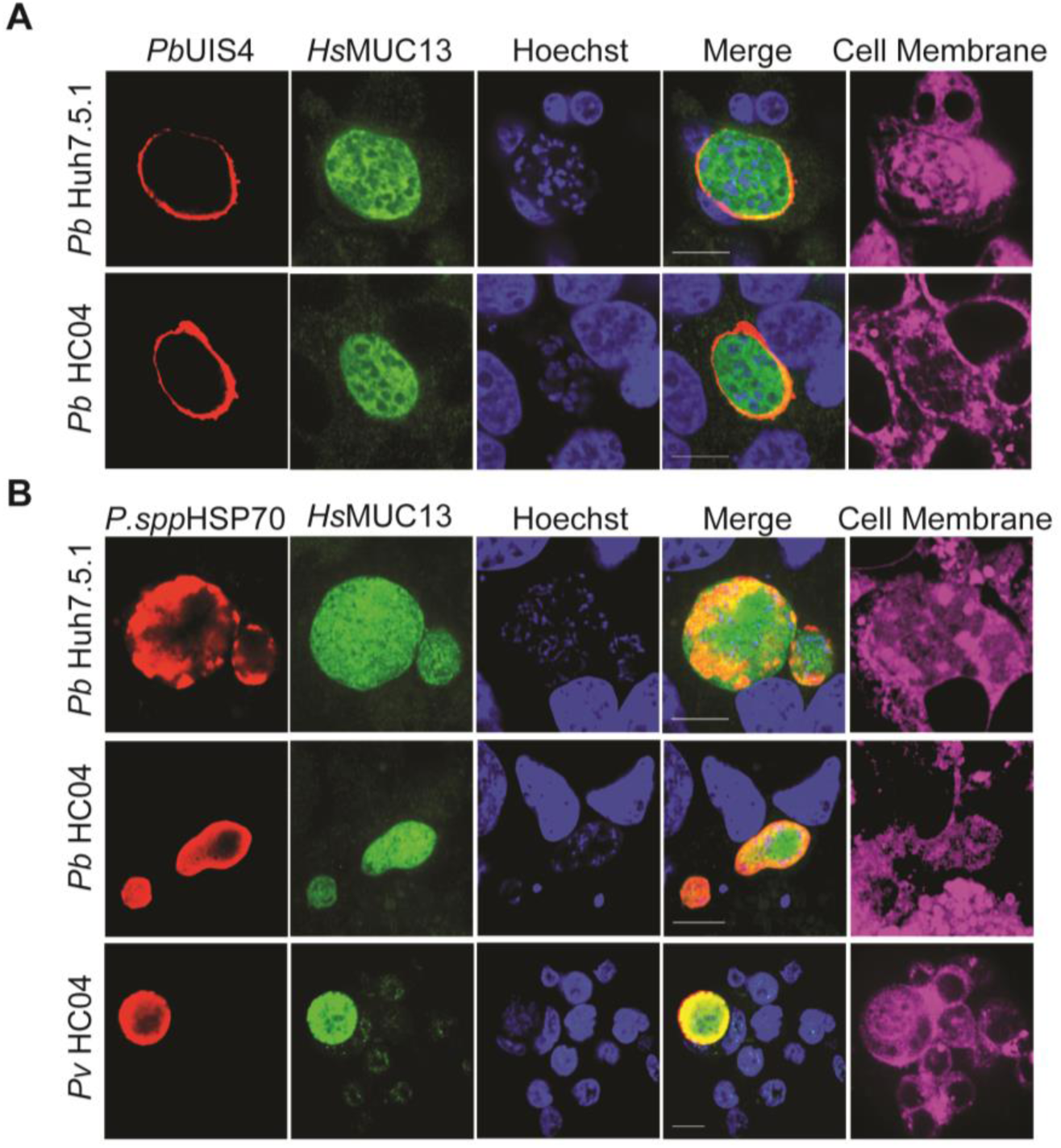
Expression and localization of *MUC13* in *Plasmodium* hepatoma cells. (A, B) Confocal microscopy images of Huh7.5.1 and HC04 liver cells infected with *Plasmodium* parasites 48 hpi. Cells were labeled using a rabbit polyclonal antibody (dilution 1:500, 1 mg/ml stock) against the intracellular region of MUC13 (MUC13 Antibody #1 - LifeSpan BioSciences #LS-C345092) and visualized with a goat anti-rabbit Alexa Fluor 488 (green); CellMask deep red was used for plasma membranes (purple). *P. berghei* (A, B) and *P. vivax* (B) EEFs were labeled using a goat polyclonal (dilution 1:200, 1 mg/ml stock) against *Pb*UIS4 (Biorbyt #orb11636) (A) or a mouse polyclonal (dilution 1:500, 1 mg/ml stock) *P.spp*HSP70 (B) antibodies and visualized with a bovine or a goat anti-goat or anti-mouse secondary antibody (Alexa Fluor 647, red), respectively. Nuclei were labeled with Hoechst 33342 (blue). Merged images between *Hs*MUC13, UIS4 or *P.spp*HSP70 and Hoechst are shown. Scale bars 10 μm; 100X oil objective.

**Figure 3:**
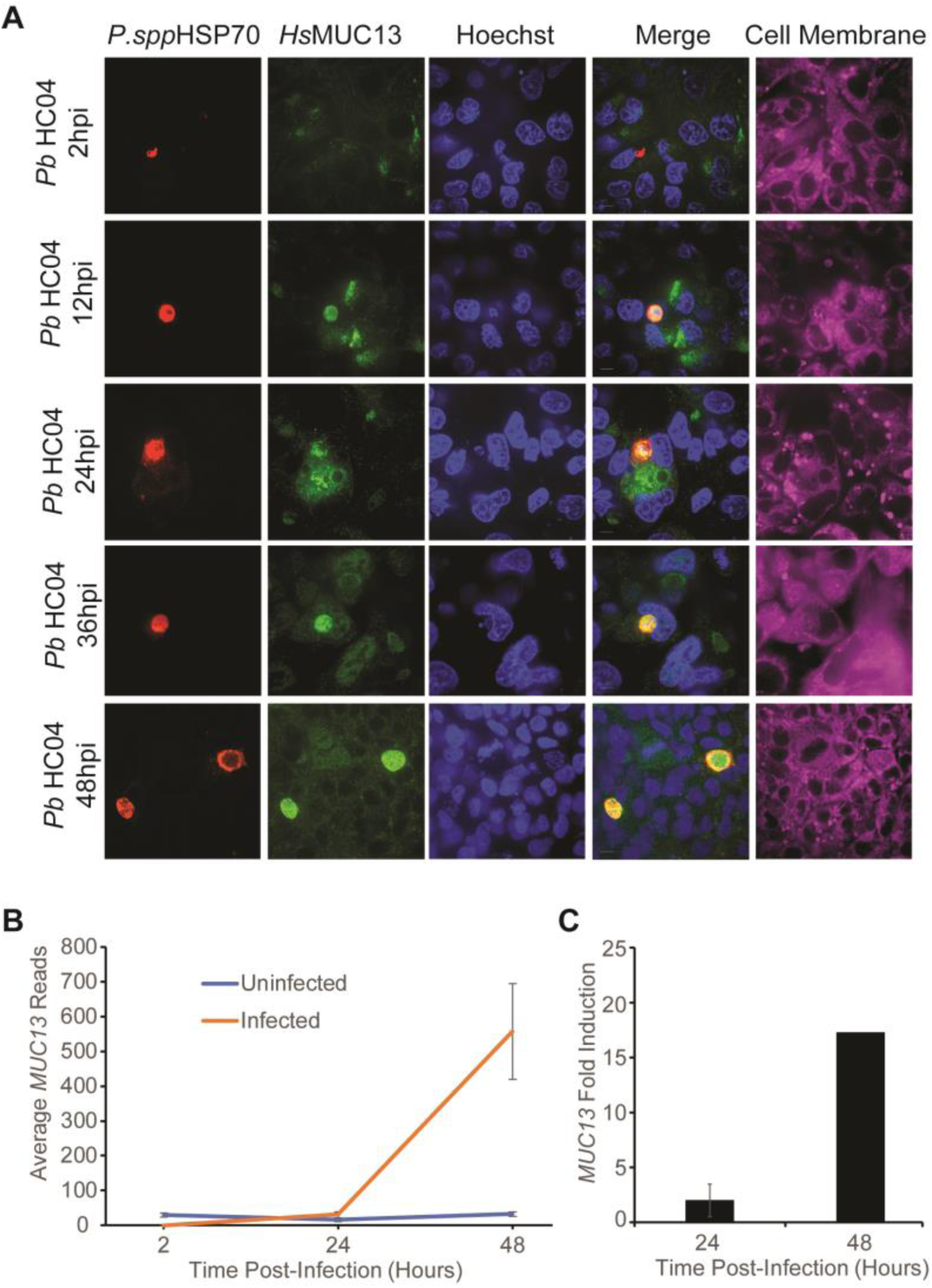
Temporal detection of MUC13 in HC04 cells infected with *P. berghei*. (A). HC04 cells were infected with *P. berghei* sporozoites and then fixed and stained at 2, 12, 24, 36, and 48 hpi. Infected cell cultures were stained using a 1:500 dilution (1 mg/ml stock) of mouse polyclonal antibody to *P.spp*HSP70 (see methods) and a 1:500 dilution of rabbit polyclonal antibody to MUC13 intracellular domain (MUC13 Antibody #1 - LifeSpan BioSciences #C345092). Primary antibody localization was visualized with goat anti-mouse (Alexa Fluor 647, red) and goat anti-rabbit (Alexa Fluor 488, green) secondary antibodies, respectively. Nuclei were stained with Hoechst 33342 (blue) and cell membranes with CellMask deep red (purple). Scale bars 10 μm; 60X oil objective. (B) The total reads, from the Huh7.5.1 RNA seq samples 1-3, for *MUC13* at the 5 indicated time points (0H Uninfected, 24H Infected, 24H Uninfected, 48H Infected and 48H Uninfected). (C) The fold-induction of *MUC13*, based upon the total read count in panel B, at 24 and 48 hpi, presented as a ratio of Infected:Uninfected.

A significant concern remained whether *MUC13* was a marker for human malaria infection, or was simply limited to the rodent model. In order to validate the utility of *MUC13* as a marker of a human malaria infection, we next examined in subcellular localization of *MUC13* in cells infected with the human parasite, *Plasmodium vivax*. Symptomatic *P. vivax* malaria patients were recruited from the Loreto region of Peru using an approved human subjects protocol and asked to provide a blood sample. This patient blood was washed and used to feed colonized female *Anopheles darlingi* mosquitos (Moreno et al., 2014) using a standard membrane-feeding assay. After 14 days, salivary glands were dissected and the resultant sporozoites were harvested and incubated with HC04 cells for seven days. Immunofluorescence microscopy demonstrated that MUC13 also colocalizes very strongly with *P. vivax* HSP70 in HC04 cells (Figure 2B), indicating that MUC13 is a marker of human malaria infection as well. Similar to the rodent parasite, MUC13 signal covers the parasite, as would be observed if *MUC13* is localized to the cytosol and nuclear membrane of developing merozoites of the developing exoerythrocytic parasite.

Species-specific parasite antibodies are typically used in liver stage imaging assays to measure parasite killing in response to drug candidates (Sinnis et al., 2013). To assess whether *MUC13* antibodies can be used as a substitute for parasite antibodies in imaging assays, we performed automated microscopy on *P. berghei* and *P. vivax* infected cultures after staining with parasite and the MUC13 antibody. *Hs*MUC13 antibodies indicated nearly identical levels of parasitemia relative to HSP70 (Figure 4A), The value of MUC13 for parasite detection was also seen in drug-response, as MUC13 was able to detect the decrease in parasite levels after dose-titration with two control compounds, atovaquone and puromycin (Figure 4B), which yielded a roughly equal EC_50_ (Figure 4C). Similar compound responses were also observed between HSP70 and *MUC13* in parasites treated at 12 and 24 hpi (Figure S6). The fluorescent signal from MUC13 antibody staining clearly demonstrates the effect of atovaquone upon parasite growth, as *P. berghei* parasites treated with atovaquone very early in the *P. berghei* lifecycle (2 hpi, when atovaquone is most effective (Swann et al., 2016)) demonstrate a marked decrease in size (Figure 4D) and infection rate when compared with later or no drug treatment. This also indicates that the MUC13 induction begins prior to 48 hpi, since parasites treated with a lethal dose of atovaquone at 12 hours post-infection still exhibit *MUC13* expression. These data show that *MUC13* can substitute as a biomarker of parasite hepatic infection.

**Figure 4:**
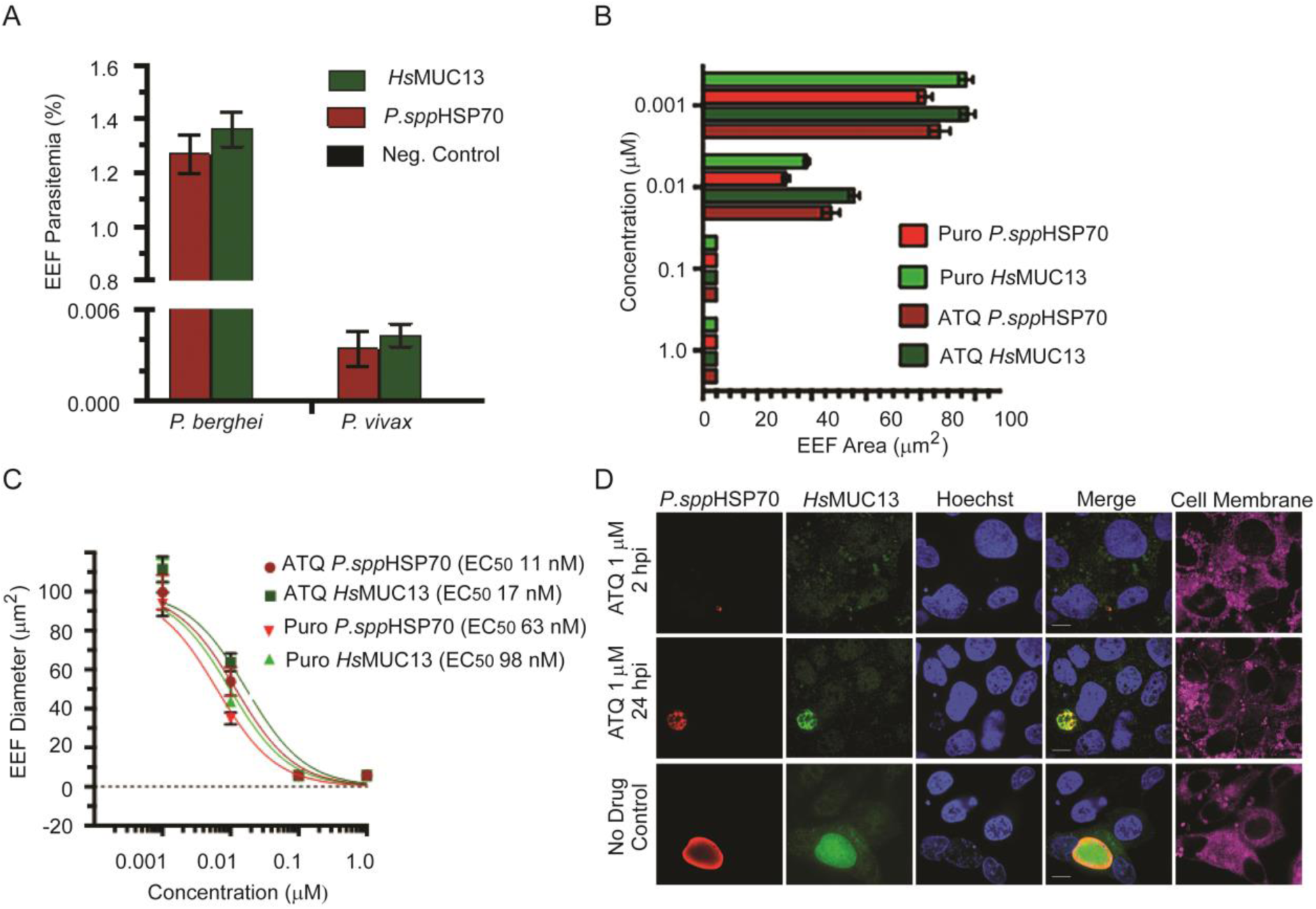
*MUC13* as a quantitative biomarker of *Plasmodium* EEF infection. (A) Detection of *P. berghei* or *P. vivax* EEF in HC04 cells by indirect immunofluorescence. Negative controls with no primary antibodies were included. Parasitemia was estimated by counting at least 240,000 cells. Three technical replicates were performed; error bars show SD. (B) Effect of atovaquone (ATQ) and puromycin (PURO) treatment (2 hpi) in cell area (growth) of *P. berghei* EEF in HC04 48 hpi. Data presented as mean ± SD with n=4. Dashed line represents the DMSO only control. (C) Dose-response curves of *P. berghei* EEF in HC04 cells for atovaquone (ATQ) and puromycin (PURO). Two technical replicates were performed; error bars show SD. 95% interval confidence for EC50s = ATQ *P.spp*HSP70, 8.98-15.82; ATQ *Hs*MUC13, 11.18-26.05; PURO *Hs*MUC13, 6.29-15.29; PURO *P.spp*HSP70, 5.01-7.92. (D) Representative images of *P. berghei* EEF in HC04 cells (48 hpi) treated (2 hpi) with 1 μM of atovaquone, puromycin, or DMSO. *P. berghei* was labeled with *P.ssp*HSP70 mouse polyclonal antibody (dilution 1:500, 1 mg/ml stock). HC04 cells were labeled with a rabbit polyclonal antibody (dilution 1:500, 1 mg/ml stock) recognizing the intracellular region of *HsMUC13* (MUC13 antibody #1 - LifeSpan BioSciences #LS-C345092). Primary antibody detection was performed with goat anti-mouse (Alexa Fluor 647, red) and goat anti-rabbit (Alexa Fluor 488, green) antibodies. Nuclei and cell membranes were stained with Hoechst 33342 (blue) and CellMask deep red (purple), respectively. Scale bar 10 μm; 100X oil.

Although the *MUC13* gene can be disrupted in mice it has been shown, nevertheless, to protect human epithelial cells against inflammation by inhibiting cellular apoptosis (Sheng et al., 2017). To determine whether MUC13 may play a role in *Plasmodium* survival and persistence, we used a traditional stable pooled shRNA approach, delivered by lentivirus, to independently knockdown human *MUC13* in both HC04 and Huh7.5.1 cells. This generated an observed 60% knockdown in MUC13 protein levels (Figure 5A). An shRNA approach was used in place of the more recent, and probably more effective, CRISPR/Cas9 approaches (Mali et al., 2013) because *Plasmodium* parasites lack the machinery for RNAi (Baum et al., 2009). Therefore, RNAi knockdown of the host-factor *MUC13* would have no potential for off-target effects upon parasite genes. Stable *MUC13* knockdown HC04 and Huh7.5.1 were established, and when infected with *P. berghei* parasites expressing luciferase, we observed a 40% decrease in parasite number in both HC04 and Huh7.5.1 cells at 48 hpi (Figure 5B). This 40% decrease in parasite number likely understates the effect of *MUC13*, given it results from a partial knockdown of *MUC13*. Imaging of parasites infecting *shMUC13* Huh7.5.1 cells by immunofluorescence (Figure 5C) indicated that the MUC13 signal recognized by the antibody, both in the parasites and in the surrounding infected cells, is markedly reduced, further validating the specificity of the MUC13 antibodies’ recognition of infecting parasites. In addition, it appears that the upregulation of MUC13 is beneficial to parasite survival and development of late stage parasites.

**Figure 5:**
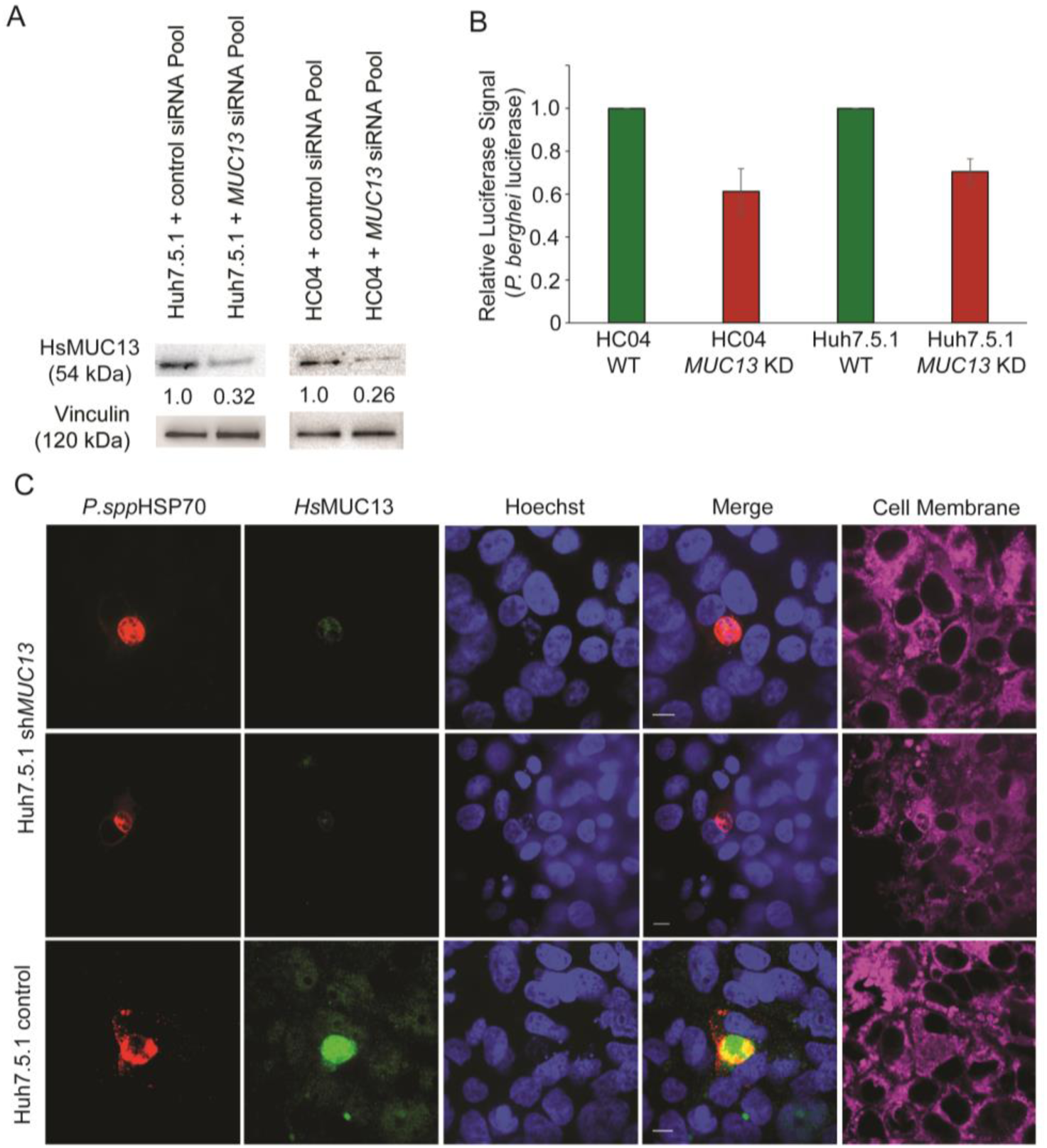
Confirmation of localization of *MUC13* in *Plasmodium* liver infected cells via shRNA knockdown. (A) Confirmation of *MUC13* knockdown in cells stably expressing an anti-mucin13 shRNA pool during infection via western blot. Values indicated are relative protein expression as determined by densitometry. (B) Effect of *MUC13* knockdown in the indicated cells infected with *P. berghei* expressing luciferase. (C) Confocal microscopy images of Huh7.5.1 liver cells, with and without *shMUC13*, infected with *P. berghei* 48 hpi. Cells were labeled with a rabbit polyclonal antibody (dilution 1:500, 1 mg/ml stock) against the intracellular domain of *Hs*MUC13 (MUC13 antibody #1 - LifeSpan BioSciences #C345092). *P. berghei* was detected using a mouse polyclonal *P.spp*HSP70 antibody (1:500 dilution, 1 mg/ml stock). Primary antibodies were detected with a goat anti-rabbit (Alexa Fluor 488, green) and a goat anti-mouse (Alexa Fluor 647, red). Cell membranes and nuclei were stained with Hoechst 33342 (blue) and CellMask deep red (purple), respectively. Merged images between *HsMUC13*, *P.spp*HSP70, and Hoechst shown. Scale bars 10 μm; 60X oil objective.

## Discussion

As the goal of malaria eradication moves forward, the need for biomarkers to identify individuals infected with malaria without clinical symptoms will be essential. One concern is the specificity of the MUC13 signal, which colocalizes with the *Plasmodium* parasite by immunofluorescence microscopy and could be due to cross-reactivity with glycolsylated parasites proteins that are only present at 24-48 hours. This is unlikely due to multiple independent lines of experimental evidence. First, the signal from the MUC13 antibody is sensitive to shRNA knockdown and shRNA knockdowns have been used by others study MUC13 phenotypes (Chauhan et al., 2012). Second, we observe no *MUC13* mRNA during early hepatocyte infection (2 hpi) or asexual blood stage infection (Figure S5), but do detect strong parasite HSP70 signal later during exoerythrocytic development. In addition, at intermediate points of infection (12 and 24 hpi), the MUC13 IFA intensity correlates with the dramatic transcriptional induction of *MUC13* during late-stage hepatic infection. In addition, the increased level of MUC13 protein within the parasite, as compared to neighboring uninfected host cells (Figure 2,3), is consistent with the 50-fold (for Huh7.5.1) induction of *MUC13* transcript levels in infected cells at 48 hpi. Third, the pattern of MUC13 expression we observe, both in host cell and parasites, is consistent with previously reported studies of *MUC13* localization (Sheng et al., 2017), displaying a speckled pattern with cytoplasmic and nuclear staining in cancer cells but apical in normal epithelial cells (Williams et al., 2001). Fourth, we find similar staining with two independently derived MUC13 antibodies that recognize different areas of the protein. Fifth, similar MUC13 localization is observed with different parasite species. Lastly, our western blots (Figure S3) do not indicate additional binding to products other than the slightly greater than 50 kDa band expected for MUC13. Our data, as well as published data, thus strongly support MUC13 specificity for both of our tested MUC13 antibodies.

How MUC13 is trafficked to the parasite cytoplasm and nucleus is an open question. Other membrane-based mucins are packaged into secretory vesicles that are targeted to the site of infection and it is possible that similar localization methods are being used here. It is possible that in other systems MUC13 is exported to the site of infection but is not observed in micrographs because this space is extracellular and protein can be washed away. Cytoplasmic and nuclear localization of MUC13 is observed in cancer cells however, where MUC13 is overexpressed (Gupta et al., 2014), so potentially a similar process of MUC13 internalization is at play here. However, in our case the protein is contained inside the parasitophorous vacuole of the parasite. Mutagenesis of the different domains of the protein will be necessary to determine which protein domains (EGF domains, SEA domain, PKC phosphorylation domain) play a role in sorting MUC13 into the parasite cytoplasm.

Under normal conditions, MUC13, like most transmembrane mucins, functions to protect cells from infection and damage by forming a barrier along the mucosal surface of those cells (Williams et al., 2001). However, data from our studies as well as those of others suggest that the cellular function of MUC13 is complex, particularly when expression is induced. Evidence suggests that MUC13 can function both as a sensor/regulatory molecule as well as an effector. Evidence that MUC13 functions as a sensor/regulator comes from overexpression and knockout studies in other systems. For example, knockout mice have an aberrant inflammatory response when challenged with pathogens (Sheng et al., 2011). In addition, MUC13 is induced in response to a variety 12 of infections, including *H. pylori* (Cheng et al., 2016). MUC13 also has been shown to interact regulatory proteins such as Her2 (Chauhan et al., 2009) and cIAP1 (Sheng et al., 2017), contains 3 EGF-like domains, which have been shown to play important immune regulatory functions in the selectins (Freedman et al., 1996), and the intracellular domain contains a protein kinase C phosphorylation domain. MUC13 is also able to move from the surface to the cytoplasm and nucleus in response to exposure to dextran sodium sulphate, a chemical shown to induce colitis (Sheng et al., 2011). Data that it also an effector molecule include the presence of mucin glycosylation domains, its upregulation after infection and, as we show here, its trafficking to the site of infection. MUC13 has also been shown to, among other functions, to increase the production of inflammatory cytokines such as IL-8 (in balance with the downregulation of cytokine production from *MUC1 (Sheng et al., 2013)*) and inhibit apoptosis (Linden et al., 2008) through the activation of Nf-kB and its resulting upregulation of downstream anti-apoptotic markers such as *Bcl-Xl, survivin* and *cyclin D1* (Sheng et al., 2017).

Since most of the transcriptional changes in host cells, such as the upregulation of cellular energy homeostasis and glucose metabolic processes and the downregulation of translational regulation and amino acid metabolism, would seem to favor parasitic development, we hypothesize that MUC13 upregulation is to the parasite’s benefit as well. In addition, the knockdown of *MUC13* appears to modestly inhibit parasite development (Figure 4). Host-cell apoptosis does represent a critical obstacle for *Plasmodium* to overcome during liver-stage infection (van de Sand et al., 2005), and inhibition of p53-mediated apoptosis specifically has been shown to be a necessary part of liver-stage parasite development (Kaushansky et al., 2013); perhaps *MUC13* induction aids late stage parasite development by delaying host cell death while the hepatocyte has detached and merozoites become fully developed (Sturm et al., 2006). There is another potential role which *MUC13* may play in parasite hepatic development, one of immune cell evasion. Given the localization of *MUC13* throughout the developing merosome, it appears that *MUC13* may be colocalized to the membranes of the developing merozoites. This could represent a parasite strategy to evade the host-cell immune system by covering the parasite membrane in host-cell glycoproteins. This strategy has been demonstrated in several other human parasites, most notably *Trypanosoma cruzi*, where the parasite surrounds itself with mucin-like glycoproteins to resemble to confuse the host immune system (Flavia Nardy et al., 2015; Nardy et al., 2016). Additionally, numerous helminths, including *Schistosoma mansoni,* express mucin-like molecules on their exterior to evade immune detection and aid invasion (reviewed in (Theodoropoulos et al., 2001)). Covering the surface of emerging merozoites with a host glycoprotein would allow those merozoites to more freely evade the immune system immediately after hepatocyte rupture and allow for more efficient erythrocyte invasion.

A final question is whether the MUC13 could be a clinically relevant biomarker of parasite infection. MUC13 protein expression has been implicated as a marker of several diseases, including numerous cancers, such as gastric, ovarian and colon cancer, inflammation and *Helicobacter* infection (Liu et al., 2014; Maher et al., 2011; Sheng et al., 2011), suggesting that MUC13 may be a general signal of cellular infection. However, these studies do not necessarily preclude the use of MUC13 as a biomarker of *Plasmodium* infection for two reasons: First, the fold-induction of MUC13 in other infectious diseases, outside of cancer, is much smaller than observed in this study (1.5-3 fold within the specific tissue assayed) (Liu et al., 2014; Sheng et al., 2011), and second, the clinical overlap between metastatic cancer patients, where MUC13 is often highly overexpressed, and malaria patients might be small. Work on MUC13 in cancer has shown that it can be detected in peripheral blood (Maher et al., 2011), suggesting that this host-factor could also potentially be used in malaria detection using peripheral blood as well. In addition, because of its role in parasite exoerythrocytic maturation, and since MUC13 is not essential gene, at least in mice (Sheng et al., 2011), MUC13 might be a target for antimalarial prophylaxis as well.

In summary, the identification of *MUC13* as a new host factor involved in parasite development will greatly enhance our understanding of parasite liver-stage development, serve as a valuable experimental window into exoerythrocytic development and potentially could serve as a clinically relevant biomarker as well.

## Acknowledgments

We thank the members of the Winzeler and Lewis labs for advice and critical reading of the manuscript. In addition, we thank Medicines for Malaria Venture for all of their support of the insectary in Peru. We would also like to thank the UCSD Institute for Genomic Medicine Sequencing Core Facility and the UCSD Human Embryonic Stem Cell Flow Cytometry Core Facility for their technical support. G.L. is supported by an A.P. Giannini Post-Doctoral Fellowship. E.A.W. is supported by grants from the NIH (5R01AI090141 and R01AI103058). N.E.L. and S.L received funding from the NIGMS (R35 GM119850) and the Novo Nordisk Foundation through the Center for Biosustainability at the Technical University of Denmark (NNF10CC1016517). The *P. vivax* work was supported by grants to J.M.V from the NIH (D43TW007120 and U19AI089681). A.N.C received support from a NIH T32 AI 007036 training grant.

### Author Contributions

G.L., P.O.S., L.W., A.C. and E.A.W. designed the experiments and wrote the manuscript. Parasite Infection, flow sorting and RNA extraction were performed by G.L., P.O.S., L.W. and J.S. Transcriptome sequencing was performed by the UCSD Institute for Genomic Medicine Core Facility. *P. vivax* experiments were designed by J.M.V, P.O.S., and M.M. and performed by P.O.S., Z.V.G., M.M. and C.T.R. Sequence analysis was performed by S.L., A.M.A. and N.L. Immunofluorescence assays were designed and performed by P.O.S and E.A.W. Knockdown of *MUC13* and subsequent determination of infection rate was performed by G.L. and B.Y.Z. All authors read and approved the manuscript. The authors declare no conflicts of interest or competing financial interests.

## References

Baum, J., Papenfuss, A.T., Mair, G.R., Janse, C.J., Vlachou, D., Waters, A.P., Cowman, A.F.,Crabb, B.S., and de Koning-Ward, T.F. (2009). Molecular genetics and comparative genomics reveal RNAi is not functional in malaria parasites. Nucleic Acids Research 37, 3788–3798.

Burrows, J.N., van Huijsduijnen, R.H., Mohrle, J.J., Oeuvray, C., and Wells, T.N. (2013). Designing the next generation of medicines for malaria control and eradication. Malaria journal 12, 187.

Chauhan, S.C., Ebeling, M.C., Maher, D.M., Koch, M.D., Watanabe, A., Aburatani, H., Lio, Y., and Jaggi, M. (2012). MUC13 Mucin Augments Pancreatic Tumorigenesis. Molecular cancer therapeutics 11, 24–33.

Chauhan, S.C., Vannatta, K., Ebeling, M.C., Vinayek, N., Watanabe, A., Pandey, K.K., Bell, M.C., Koch, M.D., Aburatani, H., Lio, Y., et al. (2009). Expression and functions of transmembrane mucin MUC13 in ovarian cancer. Cancer research 69, 765–774.

Cheng, L., Mirko, R., Sara, L., Medea, P., Caroline, B., Eva, B., Myrthe, J., Bram, F., Wim, V.D., Richard, D., et al. (2016). The Helicobacter heilmannii hofE and hofF Genes are Essential for Colonization of the Gastric Mucosa and Play a Role in IL-1beta-Induced Gastric MUC13 Expression. Helicobacter 21, 504–522.

Cloney, R. (2016). Microbial genetics: Dual RNA-seq for host-pathogen transcriptomics. Nature Reviews Genetics 17, 126–127.

Dillon, L.A., Suresh, R., Okrah, K., Corrada Bravo, H., Mosser, D.M., and El-Sayed, N.M. (2015). Simultaneous transcriptional profiling of Leishmania major and its murine macrophage host cell reveals insights into host-pathogen interactions. BMC Genomics 16, 1108.

A., Flavia Nardy, Freire-de-Lima, Geraldo, C., and Morrot, A. (2015). Immune Evasion Strategies of Trypanosoma cruzi. Journal of Immunology Research 2015, 7.

Franke-Fayard, B., Trueman, H., Ramesar, J., Mendoza, J., van der Keur, M., van der Linden, R., Sinden, R.E., Waters, A.P., and Janse, C.J. (2004). A Plasmodium berghei reference line that constitutively expresses GFP at a high level throughout the complete life cycle. Mol Biochem Parasitol 137, 23–33.

Freedman, S.J., Sanford, D.G., Bachovchin, W.W., Furie, B.C., Baleja, J.D., and Furie, B.(1996). Structure and function of the epidermal growth factor domain of P-selectin. Biochemistry 35, 13733–13744.

Gazzinelli, R.T., Kalantari, P., Fitzgerald, K.A., and Golenbock, D.T. (2014). Innate sensing of malaria parasites. Nature reviews Immunology 14, 744–757.

Gupta, B.K., Maher, D.M., Ebeling, M.C., Stephenson, P.D., Puumala, S.E., Koch, M.R., Aburatani, H., Jaggi, M., and Chauhan, S.C. (2014). Functions and regulation of MUC13 mucin in colon cancer cells. Journal of gastroenterology 49, 1378–1391.

Gupta, B.K., Maher, D.M., Ebeling, M.C., Sundram, V., Koch, M.D., Lynch, D.W., Bohlmeyer, T., Watanabe, A., Aburatani, H., Puumala, S.E., et al. (2012). Increased expression and aberrant localization of mucin 13 in metastatic colon cancer. The journal of histochemistry and cytochemistry: official journal of the Histochemistry Society 60, 822–831.

Hanson, K.K., Ressurreigäo, A.S., Buchholz, K., Prudêncio, M., Herman-Ornelas, J.D., Rebelo, M., Beatty, W.L., Wirth, D.F., Hänscheid, T., Moreira, R., et al. (2013). Torins are potent antimalarials that block replenishment of Plasmodium liver stage parasitophorous vacuole membrane proteins. Proceedings of the National Academy of Sciences of the United States of America 110, E2838–E2847.

Ingmundson, A., Alano, P., Matuschewski, K., and Silvestrini, F. (2014). Feeling at home from arrival to departure: protein export and host cell remodelling during Plasmodium liver stage and gametocyte maturation. Cellular microbiology 16, 324–333.

Ingmundson, A., Nahar, C., Brinkmann, V., Lehmann, M.J., and Matuschewski, K. (2012). The exported Plasmodium berghei protein IBIS1 delineates membranous structures in infected red blood cells. Molecular microbiology 83, 1229–1243.

Jensen, K.D., Hu, K., Whitmarsh, R.J., Hassan, M.A., Julien, L., Lu, D., Chen, L., Hunter, C.A., and Saeij, J.P. (2013). Toxoplasma gondii rhoptry 16 kinase promotes host resistance to oral infection and intestinal inflammation only in the context of the dense granule protein GRA15. Infect Immun 81, 2156–2167.

Kaushansky, A., Douglass, A.N., Arang, N., Vigdorovich, V., Dambrauskas, N., Kain, H. S., Austin, L.S., Sather, D.N., and Kappe, S.H.I. (2015). Malaria parasites target the hepatocyte receptor EphA2 for successful host infection. Science (New York, NY) 350, 1089–1092.

Kaushansky, A., Ye, A.S., Austin, L.S., Mikolajczak, S.A., Vaughan, A.M., Camargo, N., Metzger, P.G., Douglass, A.N., MacBeath, G., and Kappe, S.H. (2013). Suppression of host p53 is critical for Plasmodium liver-stage infection. Cell Rep 3, 630–637.

Kramer, A., Green, J., Pollard, J., Jr., and Tugendreich, S. (2014). Causal analysis approaches in Ingenuity Pathway Analysis. Bioinformatics (Oxford, England) 30, 523–530.

Le Roch, K.G., Zhou, Y., Blair, P.L., Grainger, M., Moch, J.K., Haynes, J.D., De La Vega P., Holder, A.A., Batalov, S., Carucci, D.J., et al. (2003). Discovery of gene function by expression profiling of the malaria parasite life cycle. Science 301, 1503–1508.

Linden, S.K., Sutton, P., Karlsson, N.G., Korolik, V., and McGuckin, M.A. (2008). Mucins in the mucosal barrier to infection. Mucosal immunology 1, 183–197.

Liu, C., Smet, A., Blaecher, C., Flahou, B., Ducatelle, R., Linden, S., and Haesebrouck, F. (2014). Gastric de novo Muc13 expression and spasmolytic polypeptide-expressing metaplasia during Helicobacter heilmannii infection. Infect Immun 82, 3227–3239.

Maher, D.M., Gupta, B.K., Nagata, S., Jaggi, M., and Chauhan, S.C. (2011). Mucin 13: structure, function, and potential roles in cancer pathogenesis. Molecular Cancer Research 9, 531 –537.

Mali, P., Yang, L., Esvelt, K.M., Aach, J., Guell, M., DiCarlo, J.E., Norville, J.E., and Church, G.M. (2013). RNA-Guided Human Genome Engineering via Cas9. Science (New York, NY) 339, 823–826.

Matuschewski, K., Ross, J., Brown, S.M., Kaiser, K., Nussenzweig, V., and Kappe, S.H. (2002). Infectivity-associated changes in the transcriptional repertoire of the malaria parasite sporozoite stage. The Journal of biological chemistry 277, 41948–41953.

Melo, M.B., Jensen, K.D., and Saeij, J.P. (2011). Toxoplasma gondii effectors are master regulators of the inflammatory response. Trends in parasitology 27, 487–495.

Melo, M.B., Nguyen, Q.P., Cordeiro, C., Hassan, M.A., Yang, N., McKell, R., Rosowski, E.E., Julien, L., Butty, V., Darde, M.L., et al. (2013). Transcriptional analysis of murine macrophages infected with different Toxoplasma strains identifies novel regulation of host signaling pathways. PLoS pathogens 9, e1003779.

Moreno, M., Tong, C., Guzman, M., Chuquiyauri, R., Llanos-Cuentas, A., Rodriguez, H., Gamboa, D., Meister, S., Winzeler, E.A., Maguina, P., et al. (2014). Infection of laboratory-colonized Anopheles darlingi mosquitoes by Plasmodium vivax. Am J Trop Med Hyg 90, 612–616.

Mueller, A.K., Camargo, N., Kaiser, K., Andorfer, C., Frevert, U., Matuschewski, K., and Kappe, S.H. (2005). Plasmodium liver stage developmental arrest by depletion of a protein at the parasite-host interface. Proc Natl Acad Sci U S A 102, 3022–3027.

Nardy, A.F.F.R., Freire-de-Lima, L., Freire-de-Lima, C.G., and Morrot, A. (2016). The Sweet Side of Immune Evasion: Role of Glycans in the Mechanisms of Cancer Progression. Frontiers in Oncology 6, 54.

Olotu, A., Fegan, G., Wambua, J., Nyangweso, G., Leach, A., Lievens, M., Kaslow, D.C., Njuguna, P., Marsh, K., and Bejon, P. (2016). Seven-year efficacy of RTS, S/AS01 malaria vaccine among young African children. New England Journal of Medicine 374, 2519–2529.

Orito, Y., Ishino, T., Iwanaga, S., Kaneko, I., Kato, T., Menard, R., Chinzei, Y., and Yuda, M. (2013). Liver-specific protein 2: a Plasmodium protein exported to the hepatocyte cytoplasm and required for merozoite formation. Molecular microbiology 87, 66–79.

Pittman, K.J., Aliota, M.T., and Knoll, L.J. (2014). Dual transcriptional profiling of mice and Toxoplasma gondii during acute and chronic infection. BMC Genomics 15, 806.

Renia, L., Mattei, D., Goma, J., Pied, S., Dubois, P., Miltgen, F., Nussler, A., Matile, H., Menegaux, F., Gentilini, M., et al. (1990). A malaria heat-shock-like determinant expressed on the infected hepatocyte surface is the target of antibody-dependent cell-mediated cytotoxic mechanisms by nonparenchymal liver cells. European journal of immunology 20, 1445–1449.

Rosowski, E.E., Lu, D., Julien, L., Rodda, L., Gaiser, R.A., Jensen, K.D., and Saeij, J.P. (2011). Strain-specific activation of the NF-kappaB pathway by GRA15, a novel Toxoplasma gondii dense granule protein. The Journal of experimental medicine 208, 195–212.

Saeij, J.P., Boyle, J.P., Coller, S., Taylor, S., Sibley, L.D., Brooke-Powell, E.T., Ajioka, J.W., and Boothroyd, J.C. (2006). Polymorphic secreted kinases are key virulence factors in toxoplasmosis. Science 314, 1780–1783.

Saeij, J.P., Coller, S., Boyle, J.P., Jerome, M.E., White, M.W., and Boothroyd, J.C. (2007). Toxoplasma co-opts host gene expression by injection of a polymorphic kinase homologue. Nature 445, 324–327.

Sheng, Y.H., He, Y., Hasnain, S.Z., Wang, R., Tong, H., Clarke, D.T., Lourie, R., Oancea, I., Wong, K.Y., Lumley, J.W., et al. (2017). MUC13 protects colorectal cancer cells from death by activating the NF-kappaB pathway and is a potential therapeutic target. Oncogene 36, 700–713.

Sheng, Y.H., Lourie, R., Linden, S.K., Jeffery, P.L., Roche, D., Tran, T.V., Png, C.W., Waterhouse, N., Sutton, P., Florin, T.H., et al. (2011). The MUC13 cell-surface mucin protects against intestinal inflammation by inhibiting epithelial cell apoptosis. Gut 60, 1661–1670.

Sheng, Y.H., Triyana, S., Wang, R., Das, I., Gerloff, K., Florin, T.H., Sutton, P., and McGuckin, M.A. (2013). MUC1 and MUC13 differentially regulate epithelial inflammation in response to inflammatory and infectious stimuli. Mucosal immunology 6, 557–568.

Singh, A.P., Buscaglia, C.A., Wang, Q., Levay, A., Nussenzweig, D.R., Walker, J.R., Winzeler, E.A., Fujii, H., Fontoura, B.M., and Nussenzweig, V. (2007). Plasmodium circumsporozoite protein promotes the development of the liver stages of the parasite. Cell 131, 492–504.

Sinnis, P., De La Vega, P., Coppi, A., Krzych, U., and Mota, M.M. (2013). Quantification of Sporozoite Invasion, Migration, and Development by Microscopy and Flow Cytometry. Methods in molecular biology (Clifton, NJ) 923, 385–400.

Sturm, A., Amino, R., van de Sand, C., Regen, T., Retzlaff, S., Rennenberg, A., Krueger, A., Pollok, J.M., Menard, R., and Heussler, V.T. (2006). Manipulation of host hepatocytes by the malaria parasite for delivery into liver sinusoids. Science 313, 1287–1290.

Swann, J., Corey, V., Scherer, C.A., Kato, N., Comer, E., Maetani, M., Antonova-Koch, Y., Reimer, C., Gagaring, K., Ibanez, M., et al. (2016). High-Throughput Luciferase-Based Assay for the Discovery of Therapeutics That Prevent Malaria. ACS Infect Dis 2, 281–293.

Tarun, A.S., Peng, X., Dumpit, R.F., Ogata, Y., Silva-Rivera, H., Camargo, N., Daly, T.M., Bergman, L.W., and Kappe, S.H.I. (2008). A combined transcriptome and proteome survey of malaria parasite liver stages. Proceedings of the National Academy of Sciences 105, 305–310.

Taylor, S., Barragan, A., Su, C., Fux, B., Fentress, S.J., Tang, K., Beatty, W.L., Hajj, H.E., Jerome, M., Behnke, M.S., et al. (2006). A secreted serine-threonine kinase determines virulence in the eukaryotic pathogen Toxoplasma gondii. Science 314, 1776–1780.

Theodoropoulos, G., Hicks, S.J., Corfield, A.P., Miller, B.G., and Carrington, S.D. (2001). The role of mucins in host-parasite interactions: Part II - helminth parasites. Trends in parasitology 17, 130–135.

Tripathi, S., Pohl, M.O., Zhou, Y., Rodriguez-Frandsen, A., Wang, G., Stein, D.A., Moulton, H.M., DeJesus, P., Che, J., Mulder, L.C.F., et al. (2015). Meta- and Orthogonal Integration of Influenza “OMICs” Data Defines a Role for UBR4 in Virus Budding. Cell host & microbe 18, 723–735.

van de Sand, C., Horstmann, S., Schmidt, A., Sturm, A., Bolte, S., Krueger, A., Lutgehetmann, M., Pollok, J.M., Libert, C., and Heussler, V.T. (2005). The liver stage of Plasmodium berghei inhibits host cell apoptosis. Molecular microbiology 58, 731–742.

Vaughan, A.M., Mikolajczak, S.A., Wilson, E.M., Grompe, M., Kaushansky, A., Camargo, N., Bial, J., Ploss, A., and Kappe, S.H. (2012). Complete Plasmodium falciparum liver-stage development in liver-chimeric mice. The Journal of clinical investigation 122, 3618–3628.

W.H.O. (2017). World Malaria Report: 2016.

Westermann, A.J., Barquist, L., and Vogel, J. (2017). Resolving host-pathogen interactions by dual RNA-seq. PLoS pathogens 13, e1006033.

Williams, S.J., Wreschner, D.H., Tran, M., Eyre, H.J., Sutherland, G.R., and McGuckin, M.A. (2001). Muc13, a novel human cell surface mucin expressed by epithelial and hemopoietic cells. The Journal of biological chemistry 276, 18327–18336.

Yalaoui, S., Zougbede, S., Charrin, S., Silvie, O., Arduise, C., Farhati, K., Boucheix, C., Mazier, D., Rubinstein, E., and Froissard, P. (2008). Hepatocyte permissiveness to Plasmodium infection is conveyed by a short and structurally conserved region of the CD81 large extracellular domain. PLoS pathogens 4, e1000010.

